# DrugDomain: the evolutionary context of drugs and small molecules bound to domains

**DOI:** 10.1101/2024.03.20.585940

**Authors:** Kirill E. Medvedev, R. Dustin Schaeffer, Nick V. Grishin

**Affiliations:** Department of Biophysics, University of Texas Southwestern Medical Center, Dallas, TX 75390; Department of Biochemistry, University of Texas Southwestern Medical Center, Dallas, TX 75390

**Keywords:** small molecules, drugs, target, domain, protein structure

## Abstract

Interactions between proteins and small organic compounds play a crucial role in regulating protein functions. These interactions can modulate various aspects of protein behavior, including enzymatic activity, signaling cascades, and structural stability. By binding to specific sites on proteins, small organic compounds can induce conformational changes, alter protein-protein interactions, or directly affect catalytic activity. Therefore, many drugs available on the market today are small molecules (72% of all approved drugs in the last five years). Proteins are composed of one or more domains: evolutionary units that convey function or fitness either singly or in concert with others. Understanding which domain(s) of the target protein binds to a drug can lead to additional opportunities for discovering novel targets. The Evolutionary Classification Of protein Domains (ECOD) classifies domains into an evolutionary hierarchy that focuses on distant homology. Previously, no structure-based protein domain classification existed that included information about both the interaction between small molecules or drugs and the structural domains of a target protein. This data is especially important for multidomain proteins and large complexes. Here, we present the DrugDomain database that reports the interaction between ECOD domains of human target proteins and DrugBank molecules and drugs. The pilot version of DrugDomain describes the interaction of 5,160 DrugBank molecules associated with 2,573 human proteins. It describes domains for all experimentally determined structures of these proteins and incorporates AlphaFold models when such structures are unavailable. The DrugDomain database is available online: http://prodata.swmed.edu/DrugDomain/

## Introduction

Proteins are vital components of cells, playing essential roles in regulating a myriad of cellular processes. The three-dimensional (3D) structure of a protein provides valuable information about protein interactions and functions. Protein domains are functionally, evolutionarily, and structurally distinct units that ensure evolutionary viability by fulfilling specific functions (Buljan and Bateman 2009; Grishin 2001). The identification and classification of protein domains into a hierarchy of evolutionary relations can lead to a better understanding of protein function by analyzing the known functions of their homologous relatives. Until recently, major structure-based classifications of protein domains have principally focused on the classification of experimentally determined protein structures: SCOP (Andreeva et al. 2020), and CATH (Sillitoe et al. 2021). Our team has developed and maintains the Evolutionary Classification Of protein Domains (ECOD), which primarily groups domains based on homology rather than topology (Cheng et al. 2014; Schaeffer et al. 2019). This feature aids in detecting instances of homology between domains with differing topologies. Another significant aspect of ECOD is its focus on distant homology, resulting in a comprehensive repository of evolutionary connections among categorized domains.

Artificial intelligence provides potent tools for scientific research across various fields, and structural computational biology is no exception. AlphaFold (AF), a recently developed deep learning method, demonstrated the capability to predict protein structure with atomic-level accuracy and has thus become an indispensable tool in structural biology (Jumper et al. 2021). Utilizing AF models, ECOD became one of the first databases to incorporate domain classification both for the entire human proteome (Schaeffer et al. 2023) and the whole proteomes of 48 model organisms (Schaeffer et al. 2024). AlphaFold has significantly expanded the available tools for computational structural biology related to drug discovery, target prediction, protein-protein and protein-ligand interaction, and prediction of complex structures (Akdel et al. 2022; Medvedev et al. 2023a; Yang et al. 2023).

Many proteins convey their functions through interaction with small organic molecules. These interactions play crucial roles in various biological processes, including enzymatic reactions, signal transduction, and regulation of gene expression. Today the majority of FDA-approved drugs in the market and their generics are small molecules (Makurvet 2021). In the last five years, small molecules accounted for 72% of all FDA-approved drugs (178 out of a total of 247 drugs) (de la Torre and Albericio 2024). Each drug has one or more protein targets with an affinity defined through experimental methods. However, many proteins are multidomain and it is commonly unknown to which domain a drug binds in most cases. Especially in the cases where the target is a large receptor. Some multidomain proteins exhibit multidomain binding sites (Kruger et al. 2012). For example, human prostaglandin D-synthase (PDB: 3EE2) binds Nocodazole (DB08313) through both domains: Thioredoxin-like (ECOD: e3ee2A2) and Repetitive alpha hairpins (ECOD: e3ee2A1) (Weber et al. 2010). Human topoisomerase II beta (PDB: 3QX3) binds anticancer drug Etoposide (DB00773) using two out of four classified domains: helix-turn-helix (ECOD: 3qx3A1) and HAD domain-like (ECOD: e3qx3A11) (Wu et al. 2011). Homologous proteins exhibit highly similar active sites, capable of accommodating similar chemical compounds (Medvedev et al. 2021; Medvedev et al. 2019). Understanding the locations and principles that govern multi-domain drug binding sites may help us to better identify them in homologous proteins and complexes made up of homologous proteins. Chemical compound repositories such as PubChem (Kim et al. 2023) and DrugBank (Wishart et al. 2018) contain information about domains of the target protein derived from Pfam (Mistry et al. 2021), which is a sequence-based classification. However, no indications are provided about to which domain(s) the compound interacts. Previously several resources were created that described ligand domain mapping using CATH and SCOP (structure-based) domain classification (Bashton et al. 2008; Bashton and Thornton 2010; Chalk et al. 2004) and Pfam database (sequence-based) (Mistry et al. 2021) classification (Kruger et al. 2012). However, these resources are out-of-date and presently unavailable for usage by the scientific community. Currently, no structure-based protein domain classification includes this type of data.

Here, we have developed the DrugDomain database, which reports those ECOD domains of proteins targeted by small molecules and drugs from DrugBank. The current version not only encompasses experimentally defined protein structures but also incorporates AlphaFold models in cases where such structures are unavailable. DrugDomain is available online at: http://prodata.swmed.edu/DrugDomain/

### DrugDomain database features and statistics

We developed the DrugDomain database (http://prodata.swmed.edu/DrugDomain/) with web interfaces that display two types of database hierarchy: protein and molecule-centric. DrugDomain catalogs ECOD domains whose residues are located within 5Å of the DrugBank molecule’s atoms. The distribution of DrugBank molecules interacting with ECOD homologous groups and architectures is shown in Figure 1. For the target proteins whose 3D structure was experimentally determined with any DrugBank molecule, the top three ECOD A-groups of the interacting domains include a+b complex topology, a/b three-layered sandwiches, and alpha arrays (Fig. 1A). The majority of small molecules interacting with a+b complex topology domains are associated with protein kinases, which are one of the most druggable protein domains and is the domain most commonly encoded among genes associated with cancer (Anderson et al. 2023; Medvedev et al. 2023b; Wang et al. 2020). The a/b three-layered sandwiches are mostly represented by Rossmann-like proteins, which were shown to bind the majority of organic molecules superclasses (Medvedev et al. 2021; Medvedev et al. 2019). For the proteins lacking their interacting DrugBank molecule in any experimental structure classified by ECOD, we predicted interacting ECOD domains using the AlphaFill algorithm (Hekkelman et al. 2023) (see below for details). In this case, the three most populated ECOD A-groups containing modeled interacting DrugBank molecules include 1) the alpha bundles with G protein-coupled receptors (GPCRs), one of the most druggable protein domains together with kinases (Wang et al. 2020), followed by 2) the a/b three-layered sandwiches and 3) a+b complex topology (Fig. 1B).

**Figure 1.**
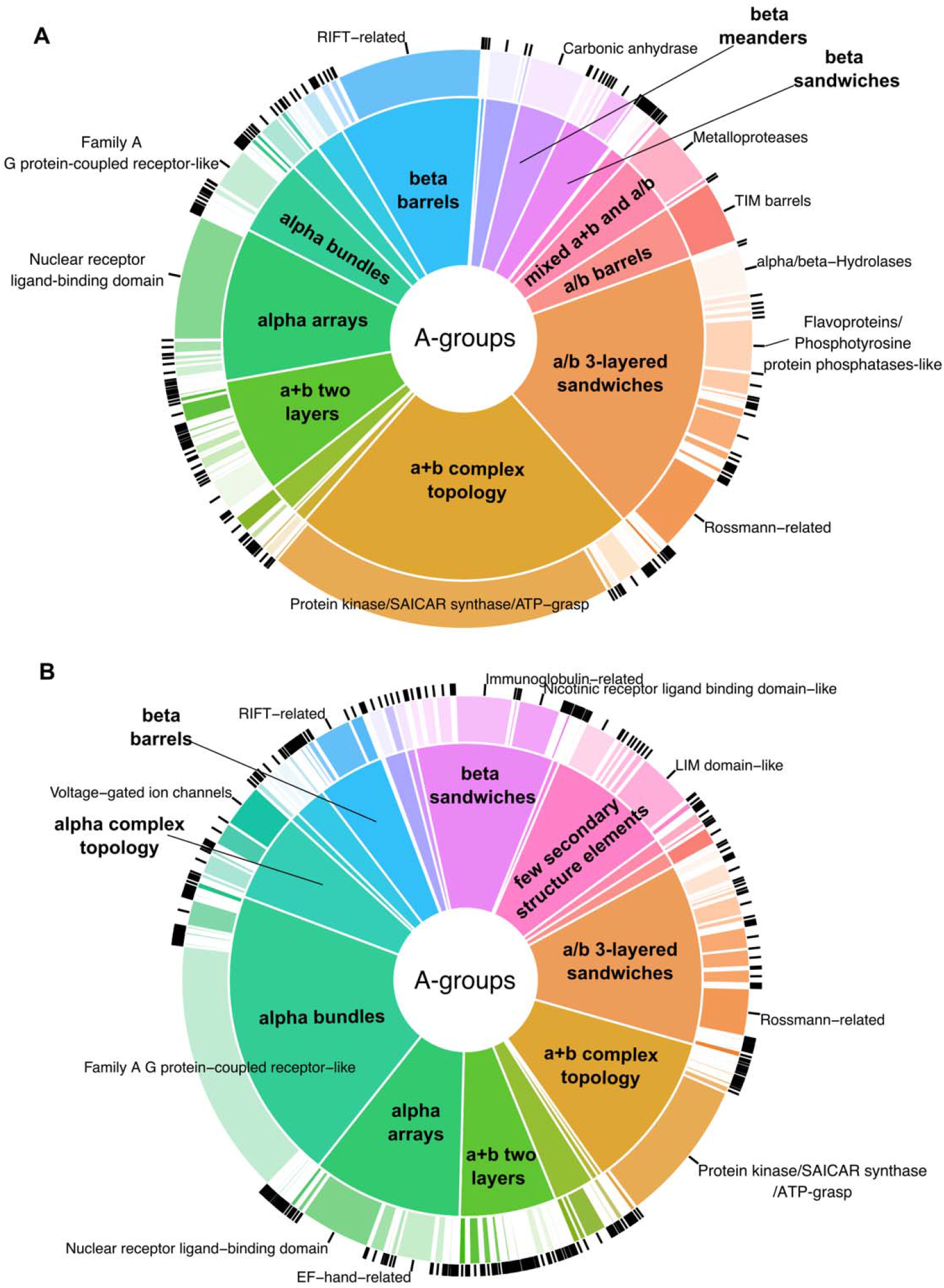
Distribution of DrugBank molecules interacting with ECOD domains of target proteins. **(A)** Distribution of ECOD domains from experimentally determined 3D protein structures determined with associated DrugBank molecule, stratified by architecture (inside pie) and homologous group (outside donut). **(B)** Similar distribution for ECOD domains with predicted small-molecule interactions using AlphaFill tool, AlphaFold models, and experimentally determined 3D structures that do not include DrugBank molecule.

The protein-centric hierarchy begins with the alphabetically sorted list of UniProt accessions (http://prodata.swmed.edu/DrugDomain/proteins/). Each accession leads to the dedicated webpage, which lists the UniProt accession, UniProt entry name, gene name, and protein name followed by the list of DrugBank molecules and drugs that target this protein. The list of molecules includes links to DrugDomain data webpages, DrugBank accession, molecule name, InChI Key, and SMILES formula. The molecule-centric hierarchy starts with the list of DrugBank accessions (http://prodata.swmed.edu/DrugDomain/molecules/). Each accession leads to the dedicated webpage, which lists the DrugBank accession, molecule name, InChI Key, and SMILES formula followed by the list of human proteins that are targets for this molecule. The list of proteins includes links to DrugDomain data webpages, UniProt accession, and protein names. The last level of our database hierarchy (DrugDomain data webpages) contains DrugBank accession, molecule name, InChI Key, and SMILES formula followed by the table of PDB structures and/or AlphaFold models with target particular protein. The table contains different information depending on the availability of experimentally determined 3D protein structures together with a particular DrugBank molecule. Such structures are available at RCSB Protein Data Bank for 2,149 molecules interacting with 783 target proteins, encompassing 5,702 PDB structures. For these cases, we identify residues whose atoms are located within 5Å of the DrugBank molecule’s atoms and map these residues to existing ECOD domains. In such instances DrugDomain data webpage’s table includes PDB accession linked to RCSB, downloadable PyMOL script, an indication that this interaction between molecule and protein target was confirmed experimentally and a list of ECOD domains interacting with the molecule with links to the ECOD database (Fig. 2A). The PyMOL script downloads the PDB structure from RCSB, colors chains by different colors, colors residues interacting with the molecule in magenta and sets their representation as sticks (Fig. 2B). We specified links to DrugDomain data webpages for corresponding domains in the ECOD database (for example: http://prodata.swmed.edu/ecod/complete/domain/e7du9A1).

**Figure 2.**
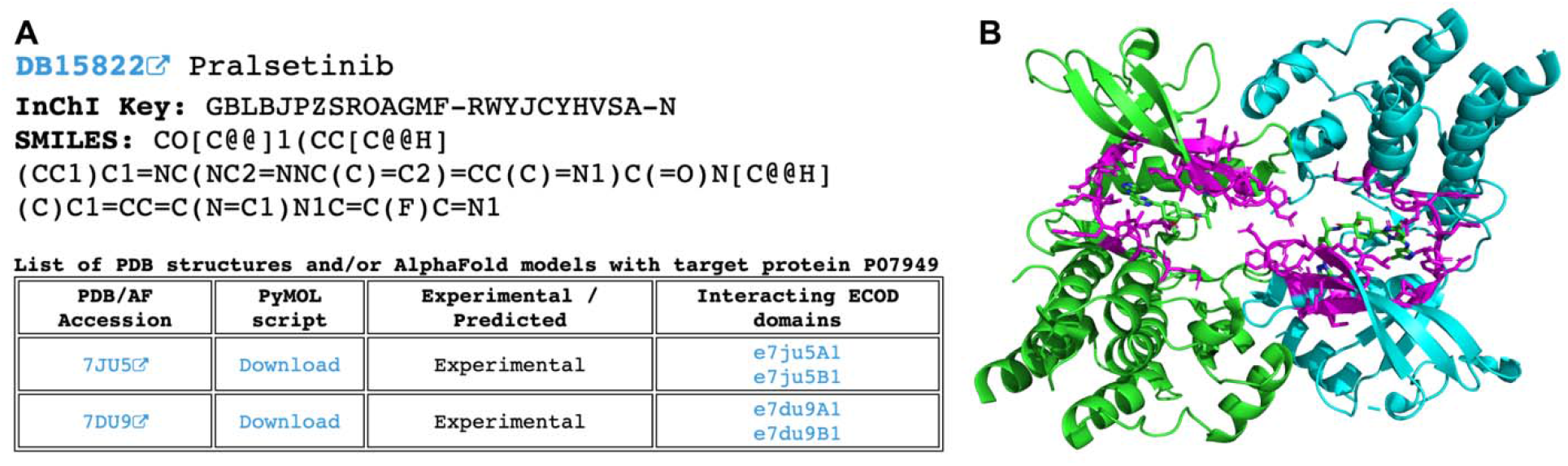
Example of the DrugDomain data webpage for cases with known PDB structure that includes DrugBank molecule and target protein. **(A)** Basic information about DrugBank molecule and table with experimental PDB structures. **(B)** Example of PyMOL script result – structure of human Proto-oncogene tyrosine-protein kinase receptor Ret (PDB: 7DU9) in complex with Pralsetinib (DB15822). Interacting residues are shown as sticks and colored in magenta.

The subset of DrugBank molecules that are not present in experimentally defined PDB structures includes 3,480 molecules targeting 2,361 human proteins. Among those proteins, experimentally defined PDB structures exist for 1,776 (75%) but without the DrugBank molecules of interest; only AlphaFold models exist for 573 (24%) proteins and there are no structural data for 12 (<1%) proteins (for example such huge proteins as E3 ubiquitin-protein ligase UBR4 (Q5T4S7), which contains 5,183 amino acids). In these cases, we applied the AlphaFill algorithm (Hekkelman et al. 2023) that uses sequence and structure similarity to retrieve small molecules and ions from experimentally determined structures to predicted protein models by AlphaFold. Using AlphaFill models we identify residues whose atoms are located within 5Å of the DrugBank molecule’s atoms of interest (if present) and map these residues to ECOD domains identified for the whole human proteome using AlphaFold models (Schaeffer et al. 2023; Schaeffer et al. 2024). If the molecule of interest is not present in the AlphaFold model, all other present ligands are considered as such. The DrugDomain data webpage’s table in these cases includes AlphaFold accession, downloadable PyMOL script, an indication that this interaction between molecule and protein target was not confirmed experimentally (and was predicted), and a list of ECOD domains interacting with the molecule with links to the AlphaFold-based ECOD database. If experimentally defined PDB structures exist for a particular protein, the table also lists them with domains from the main ECOD database (with experimental structures), which correspond to domains from the AlphaFold model. In these instances, PyMOL script is not provided due to the absence of the molecule of interest in the PDB structure (Fig. 3A). The PyMOL script includes the AlphaFold model, colors it by rainbow, colors residues interacting with the molecule in magenta and sets their representation as sticks (Fig. 3B). Full information about ECOD domains and residues interacting with DrugBank molecules are also available for download in a plain text format.

**Figure 3.**
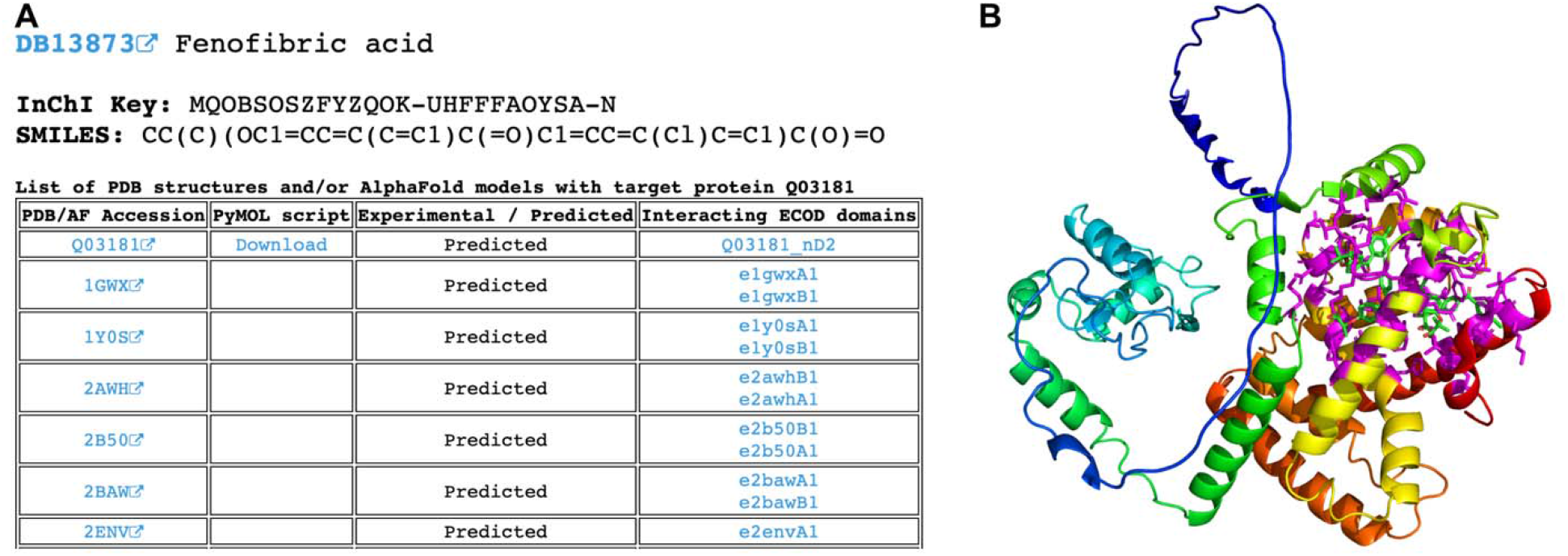
Example of the DrugDomain data webpage for cases without experimental PDB structure that includes DrugBank molecule and target protein. **(A)** Basic information about DrugBank molecule and table with AlphaFold model and corresponding PDB structures. **(B)** Example of PyMOL script result – AlphaFold model of human Peroxisome proliferator-activated receptor delta (Q03181) in complex with molecules of fenofibric acid (DB13873). Interacting residues are shown as sticks and colored in magenta.

### Collection of DrugBank molecules dataset

To obtain all drugs and small molecules that target human proteins we retrieved all DrugBank accessions (Wishart et al. 2018) related to proteins from the reference human proteome (UP000005640) using UniProt KB (UniProt 2019). Overall, using this approach we obtained 6,506 DrugBank molecules (approximately 30% of DrugBank) that are associated with 3,193 human proteins. The DrugDomain database focuses on small molecules, so we have excluded 484 DrugBank entities representing biologics (“biotech” type of molecules in DrugBank). Additionally, 224 molecules were removed from the dataset due to incomplete DrugBank records, for example, birch bark extract (DrugBank accession: DB16536) and KW-6356 (DB17080). Each molecule can have multiple targets, enzymes, transporters, and carriers associated with it in DrugBank. The purpose of this study is to catalog the interaction between DrugBank small molecules and the structural domain(s) of their target protein. Moreover, as we focus on targets that are human proteins, some molecules do not have targets in our dataset. For example, antibiotics that target bacterial proteins are excluded from the DrugDomain database (Fig. 4). Some molecules include protein groups as targets, which are often represented by various receptors. For example, Amoxapine (DB00543) targets, among others, the GABA receptor, which consists of 16 subunits, each of which is a different protein. In the majority of cases, drug targets include a particular receptor subunit known to bind the drug. However, in some cases, only protein groups are present as drug targets (for example for Mephentermine (DB01365)) and we removed such molecules from our current dataset. Thus, the remaining set, which was used for the development of the DrugDomain database version 1.0, includes 5,160 DrugBank molecules associated with 2,573 human proteins (Fig. 4).

**Figure 4.**
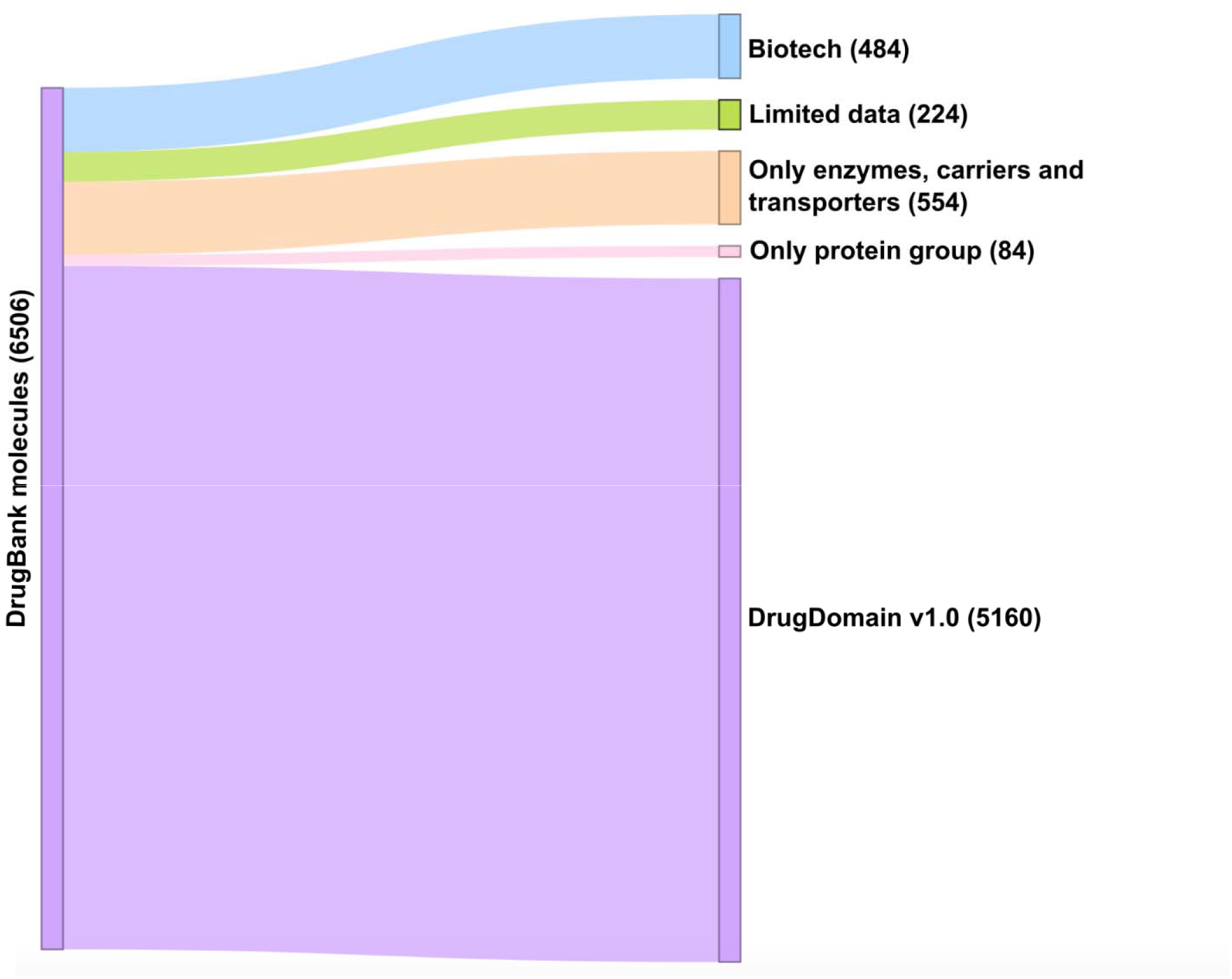
DrugBank molecules statistics for categories excluded and included in the DrugDomain database. The number of molecules for each category is shown in parentheses.

### Future perspectives

The DrugDomain database version 1.0 represents the initial step of its development. We plan to significantly expand the range of represented molecules and include all DrugBank entities in our database. Additionally, we are going to incorporate not only the protein targets of molecules but also enzymes, transporters, and carriers associated with these molecules. Considering the conservation rate of residue positions will enhance the accuracy of predicting amino acids that interact with small molecules. Finally, based on ECOD evolutionary classification and homologous evidence between proteins we anticipate to suggest new potential targets for known drugs. By focusing on evolutionarily conserved domains, we can prioritize targets that are likely to be functionally essential. Homologous proteins exhibit highly similar active sites, capable of accommodating similar chemical compounds. Better understanding along these lines opens up opportunities for discovering novel targets.

## CRediT Author Contribution

**Kirill E. Medvedev:** Conceptualization, Methodology, Software, Validation, Formal analysis, Investigation, Data Curation, Visualization, Writing - Original Draft, Project administration. **R. Dustin Schaeffer**: Software, Writing - Review & Editing, Funding acquisition. **Nick V. Grishin:** Conceptualization, Resources, Funding acquisition, Writing - Review & Editing.

